# Beyond Stimulus Onset: Ongoing Fixations Within an Object Do Not Re-evoke Category Representations During Free-Viewing

**DOI:** 10.1101/2024.12.02.623991

**Authors:** Carmel Ruth Auerbach-Asch, Leon Y Deouell

## Abstract

Human visual perception in natural conditions involves multiple fixations within single objects. While traditional studies focus on transient neural responses to initial stimuli, this study investigates how object-category representations evolve across sequential fixations on an object. Using electroencephalography (EEG) and eye-tracking, we analyzed fixation-related potentials (FRPs) and applied multivariate pattern analysis (MVPA) to decode neural representations of faces and watches during prolonged viewing. Results revealed robust category-selective responses, including the N170 component, at stimulus onset, with sustained representations persisting throughout object presentation. Temporal signal deconvolution showed that subsequent fixations did not re-evoke the N170 but elicited transient occipital responses, likely reflecting low-level differences. These findings underscore the dynamic interplay between transient and sustained neural processes during naturalistic vision and highlight the importance of disentangling overlapping neural signals during free viewing.

## Introduction

When observing natural scenes, humans tend to direct their gaze toward objects rather than the surrounding background (Buswell, 1935; Yarbus, 1967; Henderson, 2003; Einhäuser et al., 2008; Nuthmann and Henderson, 2010). This phenomenon is facilitated by humans’ remarkable capability to rapidly and effortlessly classify, recognize, and identify objects despite the computational complexity involved (Logothetis and Sheinberg, 1996; Ullman, 1996; Pinto et al., 2008). Central to object perception is categorization, where objects are naturally grouped based on shared features or properties (Rosch et al., 1976; Feldman, 1999) (Jonides and Gleitman, 1976; Olsson and Poom, 2005; Harnad, 2017). More research employing magnetoencephalography (MEG) and electroencephalography (EEG) has elucidated the temporal dynamic of category-specific representations in the brain (Rossion and Jacques, 2011; Carlson et al., 2013; van de Nieuwenhuijzen et al., 2013; Contini et al., 2017). A particularly prominent marker of category-selective activity is the N170 face-specific response, peaking at approximately 170 ms post-stimulus presentation, which signifies a distinct response evoked by faces compared to other object categories (Bentin et al., 1996; Rossion et al., 2000). Decoding studies have further revealed that basic-level category representations (e.g. snakes vs cars) begin to differentiate as early as ∼100 ms after stimulus presentation, with information pertaining to superordinate categories (e.g. animate vs inanimate) discernible around ∼240 ms post-stimulus onset (Carlson et al., 2013; van de Nieuwenhuijzen et al., 2013). However, much of this research relies on an artificial passive-viewing paradigm, where eye movements are constrained, and visual stimuli are abruptly presented at the center of the visual field. Such conditions are highly unnatural, overlooking the crucial role of self-driven eye movements in shaping perception.

In addressing this limitation, fixation-related potentials (FRPs) offer a powerful tool for studying visual responses during more natural, active vision by time-locking EEG to fixation onset rather than stimulus onset (Dimigen et al., 2011; Nikolaev et al., 2016). Our recent work, (Auerbach-Asch et al., 2020), compared FRPs elicited by self-driven eye movements (free-viewing condition) to those evoked by abrupt stimuli (fixed-viewing condition), revealing similarities in the spatiotemporal distribution of category information. In that study, parafoveal category information was hindered before the initial fixation on an object, thereby concentrating on the transient response during the first instance in which category information is available, comparable to the situation in typical passive viewing experiments. However, in natural viewing conditions, humans tend to apply multiple fixations on the same object in a seemingly redundant manner and exhibit a preference for sustaining overt attention within an object rather than immediately shifting attention to another object (Yarbus, 1967; Theeuwes et al., 2010). This propensity for multiple fixations within objects quantified as the total dwell time on an object across fixations, is beneficial for successful memory encoding (Hollingworth and Henderson, 2002), scene recognition (Nelson and Loftus, 1980; Tatler et al., 2005), and target recollection tasks (Pertzov et al., 2009; Meghanathan et al., 2019). Nonetheless, a gap remains in understanding how object representations evolve over subsequent fixations while dwelling within an object.

Recent research using deconvolution of FRPs has investigated whether fixational eye movement (particularly, smicrosaccades) during fixed-viewing on faces evoke emotion-processing effects comparable to those elicited by stimulus onset (Spiering and Dimigen, 2024). While stimulus onset elicited face-emotion-related activity, evidenced by a larger Early Posterior Negativity (EPN) for happy and angry faces compared to neutral faces, microsaccades within the face did not evoke additional emotion-related effects. Two key aspects of this study warrant further investigation to understand cognitive processing dynamics across multiple fixations. First, microsaccades during fixed-viewing tasks are relatively small (∼1.5 visual degrees), resulting in minimal retinal input changes, particularly in parafoveal vision where acuity is low and neural populations have large spatial receptive fields. The absence of fixation-related higher-level face processing effects, such as the EPN, may result from insufficient input change within the face emotion-processing machinery. Second, face emotion processing may occur later as a secondary stage of object categorization, raising the question of whether categorization itself is re-evoked during subsequent fixations.

In the present study, we investigate how object representations are maintained over subsequent fixations within a single object. Using large images and an exemplar memorization task, the task encouraged participant to make large saccades (∼6 visual degrees), which produce significant retinal changes across foveal and parafoveal neural populations. After deconvolution of overlapping E\FRPs (Dimigen et al., 2011; Smith and Kutas, 2015a; Ehinger and Dimigen, 2019), we compare the N170 face-selective component evoked by stimulus onset to that evoked by subsequent fixations within it. Furthermore, we apply multivariate pattern analysis (MVPA; Kriegeskorte and Bandettini, 2007) to the deconvolved E\FRPs to decode object categories from sequential fixations and investigate whether category representations remain stable across fixations.

## Methods

### Participants

Fifteen healthy adults (10 females and 5 males ages 18-40) with no reported neurological illness and normal or corrected to normal visual acuity, participated in the experiment. Informed consent was obtained from all participants, and they received either payment (∼$12 per hour) or class credit for their participation. The institutional ethics committee of the Hebrew University of Jerusalem approved this study.

### Experimental Apparatus and Stimuli

Participants were seated 100 cm from a 29 × 22 cm screen (ViewSonic G75f Cathode Ray Tube), set to a resolution of 1024 × 768 pixels with a 100 Hz refresh rate. The stimuli comprised 100 grayscale images of faces and 100 grayscale images of watches, all equated for luminance (mean pixel intensity) and contrast (Michelson contrast; (Lmax − Lmin)/(Lmax + Lmin), where L is luminance). Each stimulus subtended approximately 14° × 12° of visual angle, occupying the entire vertical dimension of the screen and ∼85% of its horizontal dimension. The remaining screen area had a uniform gray background (RGB = [128, 128, 128]; Figure 1). Large stimuli were chosen to encourage substantial saccades and significant retinal shifts during viewing (Figure 1). Note that the face illustrations shown in the figure are AI-generated to resemble the actual stimuli used in the experiment, which are available upon request.

**Figure 1.**
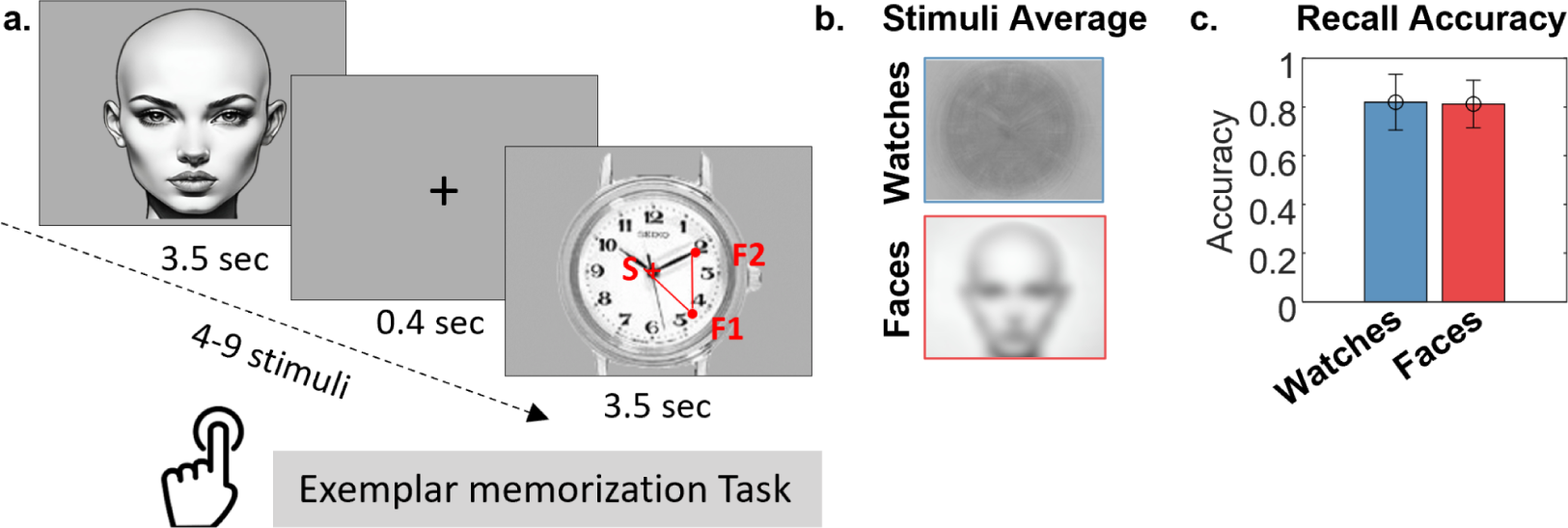
**a.** Experimental procedure: Participants memorized sequences of 4–9 stimuli (faces or watches), presented sequentially for 3.5 seconds each, and judged whether a probe stimulus was part of the sequence. Stimuli were randomly selected from a local database (the face illustrations shown are AI-generated to resemble the stimuli used in the experiment, which are available upon request). Each stimulus was preceded by a central fixation cross displayed for 400 ms. Red circles on the watch stimulus depict the sequence of events during a trial: “S” denotes stimulus onset, and “F1,” “F2,” etc., mark successive fixations. **b**. Pixel-wise averages of all watch stimuli used in the experiment and an AI-generated illustration of the average face stimuli. **c.** Mean accuracy of the two-alternative forced-choice (2AFC) stimulus recognition test across participants, with error bars representing standard deviation.

### Experimental Procedure

The experiment comprised two paradigms: Center and Off-Center. Their order was counterbalanced across participants. In the center paradigm, a fixation cross at the screen center preceded a centrally aligned stimulus (Figure 1a). In the Off-Center paradigm, the location of the fixation cross alternated on each trial between the bottom-right and bottom-left corners of the screen, and stimuli appeared parafoveally in the contralateral side of the screen (occupying either the left 85% of the screen or the right 85% of the screen respectively). Participants were required to maintain stable fixation on the cross for 400 ms before stimulus onset. Trials were aborted if they did not meet the specified criteria at the beginning of each trial. Due to instability in recording the eye position in the far left and right bottom corners of the screen, numerous trials were discarded, resulting in an insufficient number of trials for the off-center condition. Consequently, only data from the center condition were analyzed and presented in this study. A block consisted of 4-9 trials, with trial images randomly selected from a pool of all face and watch images, without replacement. These stimuli were presented sequentially, with each stimulus displayed for 3 seconds. To promote attention and exploration of the images, following each block, participants were shown an object (either a face or a watch) and asked to determine whether it had been presented in the preceding group of 4-9 stimuli. All 200 images were presented to every participant over approximately 70 blocks.

### EEG and Eye-Tracking Coregistration

EEG was acquired using an ActiveTwo system (Biosemi, The Netherlands) with 64 active electrodes mounted on an elastic cap according to the extended 10–20 system. The EEG sampling rate was 1024 Hz and an online low-pass filter with a cutoff of 1/5 of the sampling rate was applied to prevent aliasing. In addition, eight active, flat electrodes were placed: two on the mastoid processes, two horizontal electrooculogram (EOG) electrodes next to the outer canthi of the left and right eyes, two vertical EOG electrodes below and above the right eye, a single EOG electrode under the left eye, and an electrode on the tip of the nose. We referenced all electrodes during recording to a common-mode signal (CMS) electrode between POz and PO3. We recorded binocular eye movements using a desktop Eyelink 1000/2K infrared video-oculography system (SR Research Ltd., Ontario, Canada) at a sampling rate of 1000 Hz. A 9-point calibration procedure, followed by a validation stage (with acceptance criteria of worst point error < 1.5◦ visual degrees and an average error < 1.0 visual degrees), was applied before each block. We performed a gaze check before every trial by ascertaining the registration of the eyes within a 3° visual degrees radius centered on the fixation point for at least 400 ms. If this requirement was not reached within four seconds, the trial was aborted and the re-calibration procedure was repeated. EEG and eye-movement synchronization were performed using triggers sent via a parallel port from the stimulation computer, which was split and recorded simultaneously by the EEG recording system and the eye tracker. In addition to the digital eye-tracker data, we also sampled analog gaze position and pupil size from the eye tracker in the Biosemi system, simultaneously with the sampling of the EEG. We used this data to ensure accurate synchronization of EEG and eye movement data but performed eye-movement analysis on the digital data, which has a better signal-to-noise ratio (SNR).

### Eye Movement Analysis

The eye tracker provided the gaze location in the horizontal and vertical dimensions separately for the two eyes. Gaze location was computed by averaging the position of the two eyes on the horizontal and vertical axes. Blinks were defined as periods of simultaneous data loss in both eyes. Blink windows were expanded to include the closest local minima before and after the blink onset and offset with a limit of 100ms window expansion. This procedure prevents false detection of blink onset as saccades. Saccades and fixations were detected using an algorithm adapted from the EEGLab toolbox (following Engbert and Mergenthaler, 2006). First eye movement data was smoothed over a 5-sample window to suppress noise. Saccades were defined as outliers in the two-dimensional eye velocity space based on a threshold of 5 standard deviations, for a minimum duration of 10 ms, and a minimum of 30 ms inter-saccade latency. Fixation onsets were defined as saccade offsets and were included in the analysis if the saccade amplitude was under 25 degrees (determined by the size of the monitor) and if the fixation duration was under 1030 ms (determined as the cutoff of the 95% percentile of all fixation durations across participants and stimuli). On average, ∼6% (SEM = 1.3%) of fixations were removed per participant due to these inclusion criteria. In a second step, we defined Fixations of Interest (FOIs) as fixations landing within the perimeter of the object in sequential order (figure 1): First fixation (F1), second fixation (F2), and any further fixations (F3). Any fixation landing outside the object perimeter was defined as a General Fixation (GF) and included in the analysis only to account for overlapping activity (see *GLM for Overlap Deconvolution*).

### EEG Preprocessing

EEG data was preprocessed using custom Matlab code, with some functionality adopted from the Fieldtrip toolbox (Oostenveld et al., 2011), EEGLab toolbox (Delorme and Makeig, 2004) and the Unfold toolbox (Ehinger and Dimigen, 2019). Based on visual inspection, electrodes that included excessive noise across the entire duration of the experiment were excluded from the analysis (three participants lost one electrode and three participants lost two electrodes). Data were referenced to the average of all remaining electrodes. The recording was stopped between blocks. To concatenate the data to one continuous recording without amplitude step artifacts that would be later smeared by filtering, we applied linear detrending within each block by subtracting a linear vector connecting the means of the block’s initial and final ten samples. We next applied a third-degree, zero-phase-shift Butterworth high-pass filter on the data with a cutoff of 0.1 Hz. We removed 50 Hz line noise with a custom notch filter designed to suppress continuous 50 Hz oscillations, typical of power-line noise, and harmonics thereof, while having less effect on more transient components (Keren et al., 2010).

We applied independent component analysis (ICA) to attenuate noise driven by eye movements and muscle activity. For the detection of eye-movement-related ICA components, training data was generated by first removing large artifacts exceeding a threshold of 400 µV, followed by filtering between 2-100 Hz (using a 4th-degree non-causal Butterworth filter), and then concatenating segments of -200 to +1000 ms around stimuli onsets and segments of -30 to +30 ms around saccade onsets. These parameters were found to be optimal for training ICA to remove ocular artifacts (Keren et al., 2010; Dimigen, 2020). We manually identified ICA components as reflecting muscle, blink, or eye-movement artifacts, based on their temporal profile, scalp topography, and power spectrum averaged around blink, saccade, or stimulus onsets. The ratio of component variance during saccades and fixations was used as an additional measure for eye-movement-related artifacts (Plöchl et al., 2012; Dimigen, 2020). The ICA component weights, learned from the training data were next applied to the original preprocessed data (to be distinguished from the ICA training data), with the elected components removed by zeroing their weights before remixing the data into channel space (mean ± SEM number of components removed per participant: 11±1). Next, time points in which activity exceeded a threshold of ±100 µV were marked as artifacts (with an additional margin of 40 ms before and after), and rare additional artifacts were marked following a visual inspection. Finally, electrodes excluded before preprocessing were recreated by mean interpolation of the neighboring electrodes, and data were downsampled to 256 Hz for further analysis.

### Combined Temporal Deconvolution and Multivariate Analysis

We analyzed the response to faces and watches using a method for combined overlap-deconvolution and MVPA, described in Chapter 2 of this thesis (Auerbach-Asch et al., 2023). Temporal deconvolution of overlapping EEG activity is imperative in the case of fixation-related activity analysis (Nikolaev et al., 2016), due to the temporal proximity of sequential eye movements. Deconvolution can be accomplished using a general linear model (GLM; (Smith and Kutas, 2015b, 2015a; Ehinger and Dimigen, 2019). This approach yields estimates, in the form of beta coefficients, for each timepoint within a defined event of interest (e.g. stimulus onset of a specific category), reflecting the independent contribution of each overlapping event to the recorded EEG. These estimates are denoted regression Fixation-Related Potentials (rFRPs) or regression Event-Related Potentials (rERPs), replacing traditional averaged Fixation-Related and Event-Related Potentials (FRPs or ERPs, respectively). We analyze category-selective activity in rERPs\rFRPs (with temporal deconvolution) as well as their respective ERPs\FRPs (without temporal deconvolution). For details of our GLM, please refer to section *GLM for overlap deconvolution* in this chapter.

The application of multivariate pattern analysis (MVPA) methods on deconvolved rERPs\rFRPs requires the reconstruction of specific data segments from the estimated mean responses. To do this, all estimated overlapping responses, excluding a single event of interest, are subtracted from the raw data, ensuring the isolation of the signal specific to that particular event (see Auerbach-Asch et al., 2023*; “Combined Temporal Signal Decomposition and MVPA Decoding”*, and *Reconstruction of deconvolved segments* bellow). MVPA methods also mandate that model learning and testing be performed on separate datasets, to avoid overfitting the model to the data (Hastie et al., 2009; Grootswagers et al., 2017; Kriegeskorte and Douglas, 2019). Therefore, it is crucial to divide the train and test datasets before applying GLM-deconvolution and subsequent event-specific decoding.

#### Data Partitioning into train and test sets

We prepared our data for a 3-fold cross-validation procedure for decoding face vs watch stimuli from the EEG data. Face and watch trials were randomized and partitioned into three sets in which the number of trials from each category was balanced within the set. For each fold, two sets were combined to form the training set (66.7% of the data) and the remaining set was used as the test set (33.3%). Trials in each set were concatenated to produce continuous data for the GLM. This process was iterated five times, employing different fold assignments to mitigate the risk of spurious decoding arising from fold assignment variability.

#### GLM for Overlap Deconvolution

The GLM, designed to account for overlapping activity from sequential events (figure 1), included nine events: Stimulus onset (Face-S, Watch-S); first fixation (Face-F1; Watch-F1); second fixation (Face-F2; Watch-F2); further fixations (Face-F3; Watch-F3); and general fixation (GF) to account for any fixations outside the regions of interest (see *Eye-Movement Data Analysis* section). To obtain the response over time, each of these nine predictors was expanded as a set of timepoints by timepoint predictors from -200 ms to 1500 ms post-stimulus onset. Sampled at 256 Hz, this time window resulted in 436 timepoint predictors for each of the nine events (see Ehinger and Dimigen, 2019, for a detailed explanation of time-resolved GLM). Finally, to account for saccade direction and amplitude effects on FRPs, each fixation predictor was accompanied by its incoming saccade-direction continuous predictor, split to its cosine and sine values, and a saccade amplitude continuous mean-centered predictor spread over a five-spline basis set (Ehinger and Dimigen, 2019). The Wilkinson notation (Wilkinson and Rogers, 1973) of our GLM design matrix is:

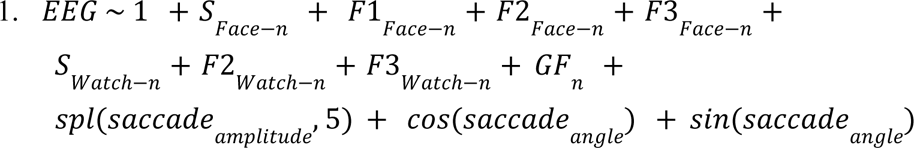

Where all predictors are binary (categorical) except those preceded by *spl* (stands for spline basis), *cos*/*sin* (stand for cosine and sine basis). The *n* under the binary predictor is a reminder that notation represents a group of predictors for each timepoint in the segment separately (Smith and Kutas, 2015b).

Finally, time points containing artifacts were excluded from the model design matrix. The GLM was solved using the Matlab lsmr function (Fong and Saunders, 2011), separately for each train and test set in each fold to avoid overfitting.

#### Reconstruction of deconvolved segments

The GLM resulting coefficients were used to reconstruct the EEG activity by summation of overlapping evoked activity from all events of interest throughout the experiment. Then raw data was segmented into time windows around the events-of-interest onsets (e.g. face \ watch onset and sequential fixations). The reconstructed overlapping activity was subtracted from each segment, excluding the contribution of the specific event of interest, resulting in deconvolved single segments. Deconvolved segments were obtained separately for each dataset in each fold, and used later for the MVPA (see chapter 2 of this thesis for more details).

#### MVPA Decoding

To quantify the representation of category-related information locked to stimulus onset (S) or ensuing fixations (F1-F3), we trained a set of linear classifiers to decode (object categories from reconstructed single presentations or fixations separately at each time point. The instantaneous voltage in each EEG-cap electrode (excluding external electrodes) created the 64-point feature vector for each time point. As noted, to avoid overfitting, we applied a 3-fold cross-validation procedure, ensuring that the GLM was calculated separately for the train and test sets (data was partitioned as explained in the *Data partitioning* section), which was repeated 5 times. To ensure model training was performed on balanced categories, the category containing more segments was undersampled to match the other category. On average, the number of events per category included in each training/testing fold was (mean ± SEM across participants): 65.5±0.4/32.7±0.2 Stimulus Onset segments; 64.9±0.4/32.3±0.3 F1 segments; 55.5±2.7/27.5±1.4 F2 segments; 355±27.9/179±14.9 F3 segments. Initially, baseline correction was applied to all segments before decoding by subtracting the mean activity between -200 to -50 ms before stimulus or fixation onset from the entire segment. Subsequently, the analysis was repeated on non-baseline-corrected trails for comparison. For classification, we applied Linear Discriminant Analysis (LDA) at each timepoint separately, using custom code based on the MVPA-Light toolbox (Treder, 2020). LDA attempts to find a linear weighting 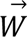 of the scalp electrodes for which separation between the categories is maximal. This is achieved by simultaneously maximizing the distance between categorical mean responses (“signal”) while minimizing response variability (“noise”), using the following formula:

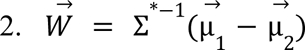

where 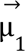 and 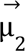 are the mean responses to each category and Σ is the covariance matrix obtained by averaging the covariance matrices of each category separately. To reduce the influence of random noise fluctuations on the weight estimate, we used a shrinkage estimator for the covariance matrix (Σ^*^; Blankertz et al., 2011) by combining the empirical covariance (Σ) and the identity matrix (*I*), weighted according to a regularization parameter (λ):

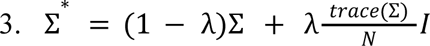

Multiplication of the identity matrix by the mean value on the covariance diagonal (N is the number of features of the classifier, equal to the number of electrodes in this case) ensures that the trace of the covariance is preserved, which helps to mitigate the bias introduced by this step. The regularization parameter (λ) was estimated using the Ledoit-Wolf formula (Ledoit and Wolf, 2004) implemented in MVPA-Light. Once the linear weights 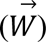 were obtained, the threshold for separating between the two categories was computed as:

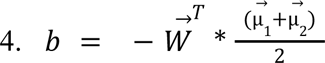

The predicted category for each test-set segment at each timepoint is determined by multiplying the vector of electrode activations 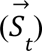 with the feature weights 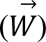 and adding the threshold term (*b*):

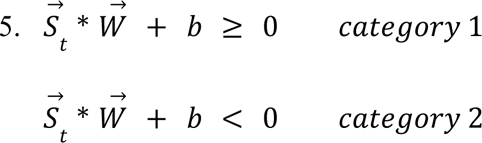

The classifier’s accuracy at each time point was calculated as the percentage of correct classifications made on the test set. This was averaged across three folds and five repetitions of the cross-validation process. Given the balanced test sets, accuracy was deemed a suitable performance metric.

For interpretability of the classifier weights in terms of activation patterns (or the extent to which each feature\electrode contributed to the classification decision), we corrected the weights using the transformation suggested by (Haufe et al., 2014) :

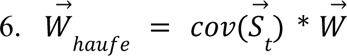

To assess the stability of representations over time, we employed the temporal generalization method (King and Dehaene, 2014; King et al., 2014). In this approach, a classifier trained on data from a specific time point was used to decode data at every other time point within the segment. Temporal Generalization Matrices (TGMs) depict decoding accuracies obtained by classifiers trained at each time point within the segment (y-axis) and tested on data from all other time points in the test segments (x-axis), with colors representing decoding accuracy levels. Stable representations entail that the classifier trained on earlier time points can accurately decode data from later time points in the segment.

### Statistical analysis

#### Behavioral Statistics

At the end of each block, which consisted of the presentation of 4-9 stimuli, participants were shown a probe stimulus and required to indicate, by button press, whether it had been presented in the preceding block or not (‘y’ for yes, ‘n’ for no). We compared the percentage of correct responses for the two stimuli categories using a paired t-test.

#### Eye Movement Statistics

Eye movement parameters, including fixation durations and saccade amplitudes, were analyzed by computing the median value for each participant for the two stimuli categories: Watches and Faces. Subsequently, a two-tailed paired-sample t-test was conducted across participants to compare these parameters between the categories with a significance threshold of p<0.05 and calculated Cohen’s effect-size value 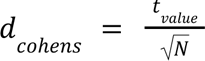 (Rosenthal and Rosnow, 2008).

#### Univariate rERP\rFRP and ERP\FRP Statistics

We compared category-specific responses after data partitioning (*Data Partitioning into train and test sets*), on the combined train and test segments of the first fold (since all folds include different partitioning of the same data). We report results without (ERPs\FRPs) and with temporal deconvolution (rERPs\rFRPs; *Reconstruction of deconvolved segments*).

Segments were low-pass filtered with a cutoff of 30 Hz using a 4th-order non-causal Butterworth filter and averaged within each participant. Initially, baseline correction was applied to all rERPs\rFRPs by subtracting the mean activity within -200 to -50 ms before event\fixation onset from the entire segment. A subsequent analysis was performed without applying baseline correction. We present the mean rERPs\rFRPs over participants with a 95% confidence interval margin around the mean.

We tested for significant differences between responses to faces and watches using a cluster permutation test (Maris and Oostenveld, 2007). For this test, a two-tailed paired t-test was conducted for each time point, and clusters were defined by temporally adjacent points that crossed a threshold of p<0.01. Points were merged into a cluster only if they were both negative or both positive. The cluster statistic was defined as the absolute sum of t-values (t_sum) of all points in the cluster. We constructed a null distribution of cluster statistics by repeating the procedure after randomly permuting the assignment of category labels (1000 permutations). From each iteration, we extracted the largest t_sum and included it in the null distribution. Finally, clusters in the real data with t_sum>95 percentile of the null distribution were considered significant (Figure 3;5).

**Figure 2:**
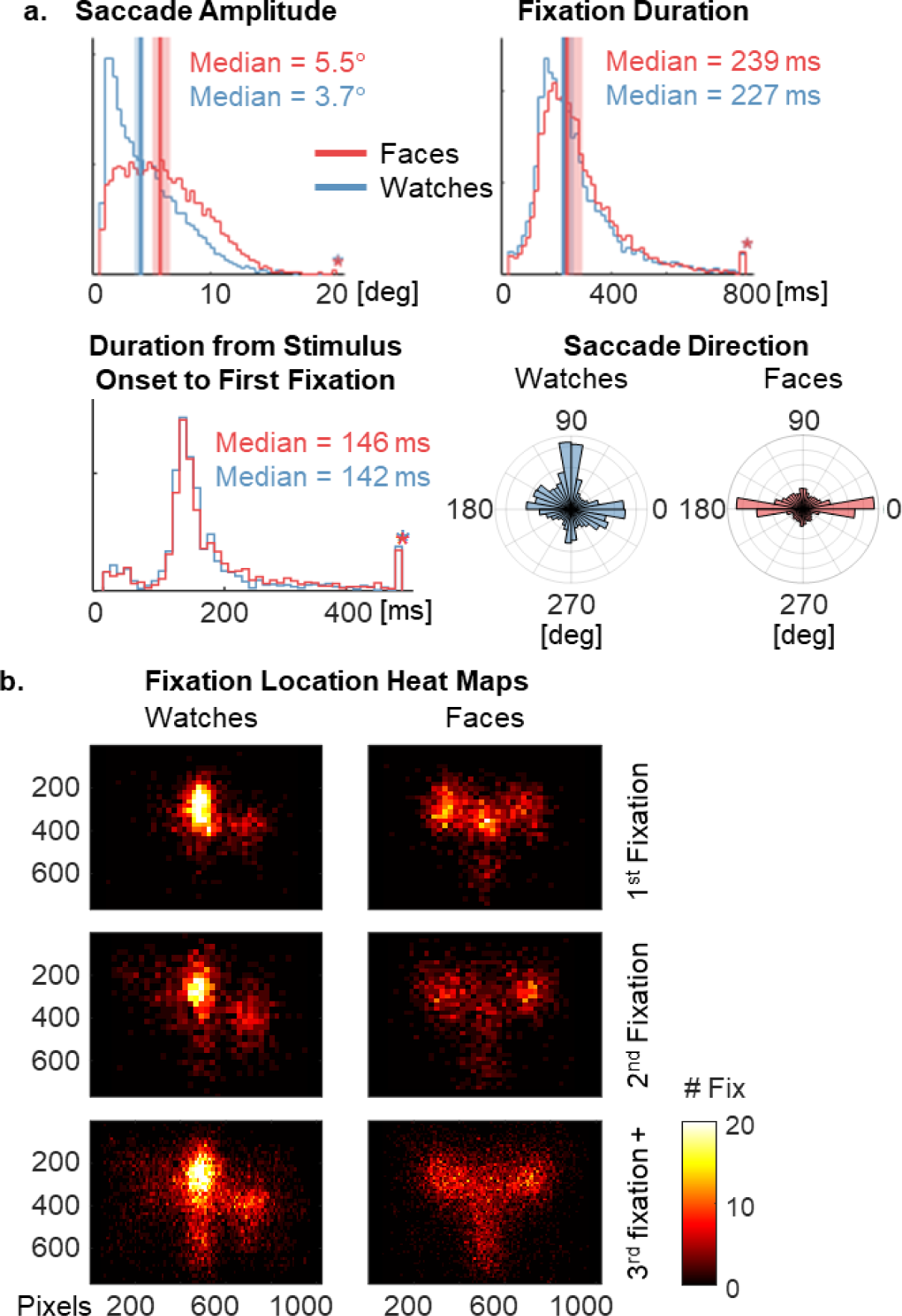
a. Histograms of eye movement parameters on watch (blue) and face (red) stimuli separately (values scaled from 0 to 1). The histograms represent all fixations included in the analysis aggregated across participants. Solid vertical lines and the shaded bars indicate the mean and standard deviation, respectively, of the single participant histogram medians. The starred column on the far right represents the bin for all values that lie off the edge of the graph (Using the function ‘Plot and compare histograms’, Lansey JC, 2024). Top left: distribution of saccade amplitudes preceding fixations. Top right: distribution of fixation durations. Bottom left: Distribution of the duration from stimulus onset to the first fixation. Bottom right: Normalized angular histograms for saccade directions. **b.** Heat maps depicting fixation locations in pixels for watch stimuli (left) and face stimuli (right) separately. The color bar stands for the number of fixations in a certain location on the screen.

**Figure 3:**
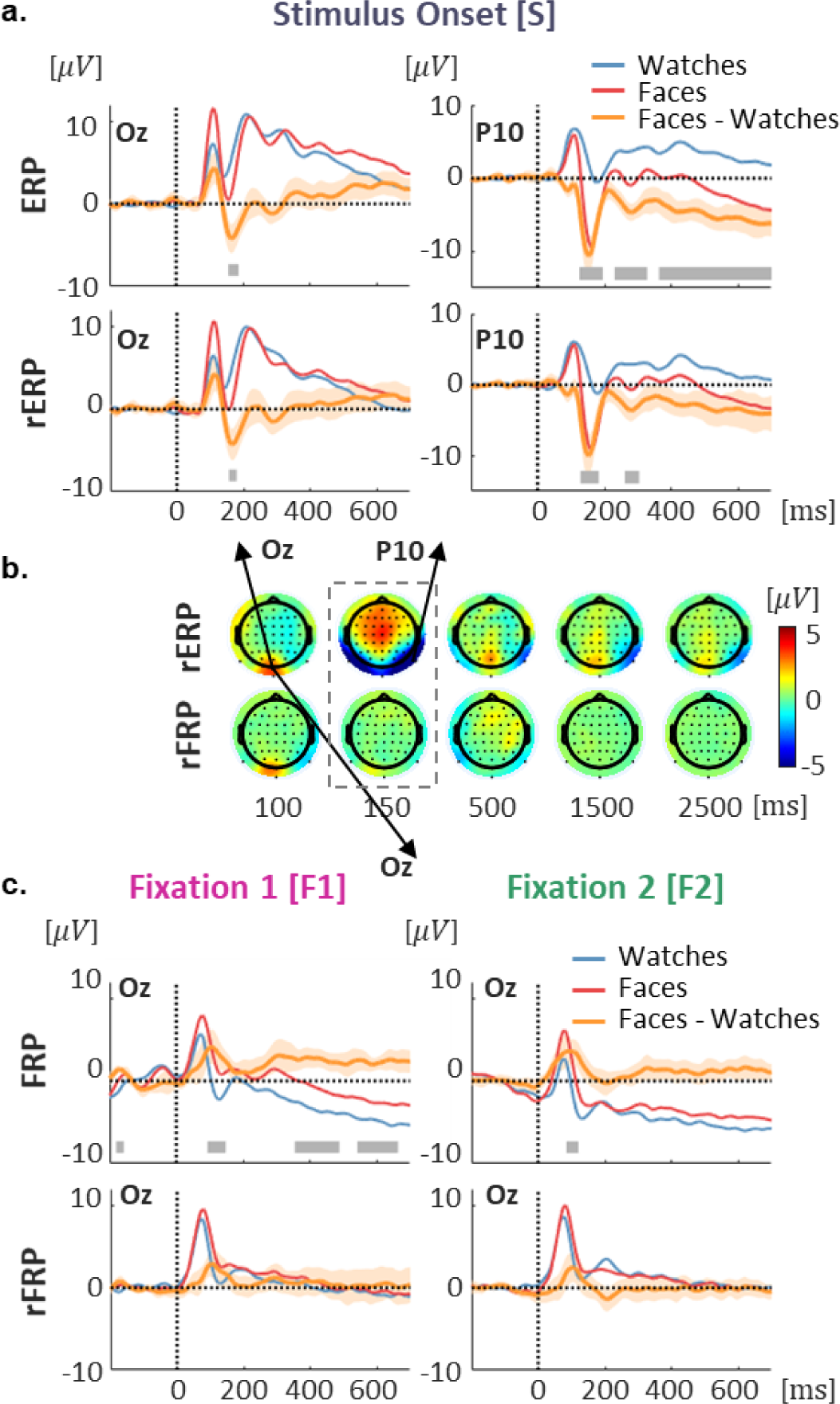
Category selective activity evoked by stimulus onset compared to ensuing fixations. a. Grand-average evoked response to watches (blue), faces (red), and the face-watch differential responses (orange), in electrodes Oz (left column) and P10 (right column), using classical segmentation and averaging (ERP; top), and deconvolution by regression (rERP; bottom). Intervals around the differential response depict 95% confidence intervals across subjects. Gray horizontal lines on the bottom depict clusters in which the differential response is significantly different than zero using a cluster permutation test. b. Scalp topography of face-watch grand-average differential response evoked by stimulus onset (top) and ensuing fixations (average of F1 and F2) after signal deconvolution. The dotted box indicates the time window associated with the N170 component. Arrows indicated the electrode locations of Oz and P10. c. Grand-average fixation-related responses in electrode Oz evoked by watches (blue), faces (red) and the face-watch differential response (orange) including 95% confidence intervals, obtained by segmentation and averaging (FRP; top) and by deconvolution by regression (rFRP; bottom).

#### Multivariate decoding Statistics

We calculated the decoding accuracy of a classifier trained to distinguish between face and watch stimuli at each timepoint within a segment (see section *MVPA Decoding*). The classification was performed on data with and without temporal deconvolution in order to evaluate the effect of deconvolution. Subsequently, we employed a cluster-based permutation test (1000 permutations) across participants based on a one-tailed one-sample t-test against a 0.5 (chance). Temporal cluster inclusion criteria were set to a threshold of p<0.05. The t-distribution under the null hypothesis was built by randomly permuting labels of the actual accuracies and chance. Finally, clusters in the real data with a t_sum value exceeding 95% of the null distribution were considered significantly different from chance. The same methodology was applied to the TGMs across participants with clusters defined by temporal adjacency in two dimensions (Figure 4;6).

**Figure 4:**
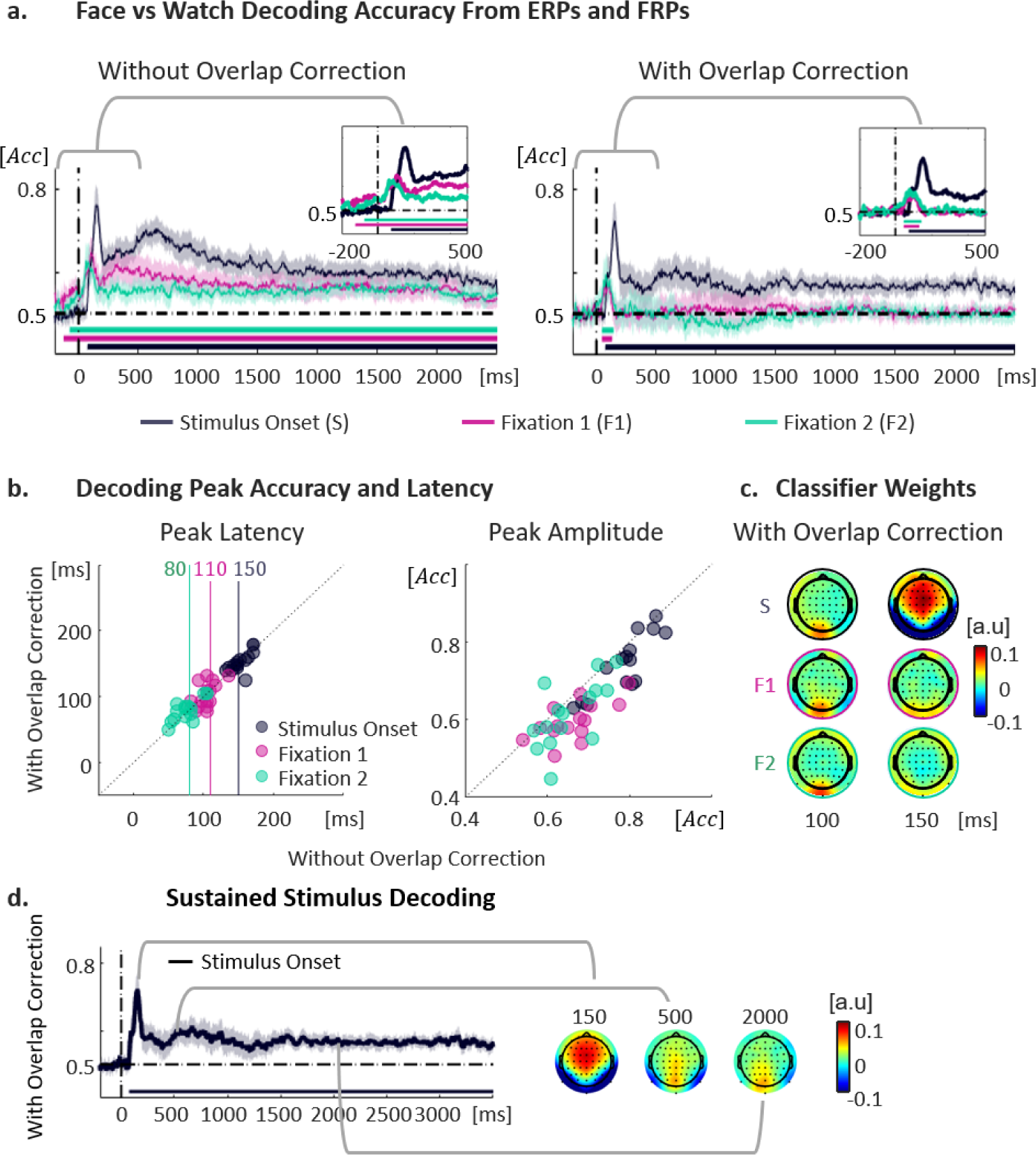
Decoding face vs. watch stimulus category from stimulus onset ERPs (S) and ensuing fixation FRPs (F1, F2). **a.** Grand average time-resolved decoding accuracy from EEG segments without (left) and with (right) temporal overlap correction, time-locked to stimulus onset (black), the first ensuing fixation within the object (F1, pink) and the second fixation (F2, green). The shaded intervals indicate 95% confidence intervals around the mean, and the horizontal bars at the bottom depict time points during which the accuracy is significantly larger than chance using a permutation test. Insets provide a zoom onto the first 500 ms (and baseline). **b.** Decoding peak latencies (left) and amplitudes (right), comparing with (y-axis) and without (x-axis) overlap correction. The solid vertical lines depict the grand-average peak latencies. The circles depict single-subject decoding peaks within ± 30 ms from the grand average. **c.** Classifier weights at 100 and 150 ms post stimulus or fixation onset, for the overlap corrected segments. **d.** The grand average time-resolved decoding accuracy from EEG segments following temporal overlap correction ranged from -200 ms to 3500 ms around stimulus onset (same data as panel a-right). The shaded intervals indicate 95% confidence intervals around the mean, and the horizontal lines depict time points during which the accuracy is significantly larger than chance using a permutation test. On the right, classifier weights are presented for 150, 500, and 2000 ms after stimulus onset.

## Results

### Behavioral results

Participants correctly determined if a probe stimulus was presented in the preceding block with an accuracy of 0. 82 ± 0. 11 (mean ± std) for watch stimuli and 0. 81 ± 0. 09 for face stimuli, with no statistical difference between categories (t(28) = 1.9, p=0.9; Figure 1).

### Eye movements

Participants executed an average of 7. 3 ± 1. 8 (mean ± std) valid fixations on watches and 7. 2 ± 1. 7 valid fixations on faces, with no statistically significant difference between the two categories (t(28) = 0.06, p=0.9). Fixations on watches were predominantly centered in the middle and top portions of the stimulus, while fixations on faces were focused on salient features such as the eyes and nose (Figure 2b). Figure 2a compares saccade and fixation parameters between the two categories. Median fixation durations across participants were not statistically different between the watch (227 ± 43 *ms*; median ± std) and face (239 ± 57 *ms*; median ± std) categories (t(28) = -1.2, p=0.2). Faces and watches also elicited similar distributions of intervals between stimulus onset and the first subsequent fixation. Taken together we expect similar temporal overlapping effects between categories. However, faces elicited larger saccade amplitudes (5. 5 ± 1. 4 *deg*; median ± std) than watches (3. 7 ± 0. 8 *deg*; median ± std; *t*(28) = − 4. 6; *p* < 0. 001; *Cohen*’*s d* = − 1. 2). In addition, saccades on faces were primarily along the horizontal axis while those on watches were slightly more frequent along the vertical axis, probably driven by the typical spatial distribution of the facial parts compared to watch parts. Both saccade amplitude and direction were accounted for in the deconvolution GLM (see *GLM for Overlap Deconvolution*), however, they may contribute to low-level processing differences.

### Do ensuing fixations evoke category-selective activity?

Traditional ERP studies of category processing typically focus on transient responses to brief presentations of stimuli (typically less than 500ms). In our study, stimuli were presented for long durations to allow participants to perform multiple fixations within the same object, and we asked if category-specific activity is re-evoked with every sequential fixation. Typically, VEPs are maximal over occipital electrodes (Oz), and category-selective responses are most prominent over occipitotemporal electrodes (P10). Figure 3a depicts the evoked response to stimulus onset in electrodes Oz and P10, obtained by standard segmentation and averaging (ERP) and by the deconvolution by regression method (rERP). Using both methods, we observed a robust stimulus-evoked N170 component, with a larger negative response for faces compared to watches, predominantly over lateral occipitotemporal electrodes (P9\P10; (Bentin et al., 1996). The differential face-watch activity peaked approximately 150 ms post-stimulus onset, accompanied by a reversed polarity Vertex Positivity Potential (VPP; (Jeffreys, 1996) over center-frontal electrodes (Figure 3b-top row). However, ensuing fixations following the stimulus onset (F1, F2) elicited an early and transient occipital response known as the FRP lambda component (Yagi, 1981; Kazai and Yagi, 1999; Rajkai et al., 2008), at approximately 100 ms post-fixation onset, but no N170 differential component (Figure 3b-c). rFRPs, isolating the fixation-related responses from the preceding stimulus-evoked response, did not differ between the face and watch categories (Figure 3c bottom) leading to the conclusion that category-selective activity indicative by the N170 component, is not re-evoked by ensuing fixations within the object.

Traditional univariate analysis, which focuses on individual electrodes or specific time windows, may lack the statistical sensitivity needed to detect visual representation differences that are distributed across space and time. This issue is particularly evident when averaging across participants, as the spatial distribution of effects can vary considerably between individuals. To address this limitation, we employed MVPA decoding of object category (faces vs watches) in individual subjects, utilizing relationships among all electrodes, and leveraging not only the mean category information but also the response variance across trials. Figure 4 presents time-resolved decoding accuracies for segments time-locked to stimulus onset versus those time-locked to subsequent fixations (F1, F2). Our results demonstrate that stimulus-evoked activity yields higher decoding accuracies than fixation-related activity, with a noticeable impact of signal deconvolution on decoding outcomes (Figure 4a). Segments time-locked to stimulus onset produce sustained above-chance decoding, an effect that is disentangled from subsequent fixations only after overlap correction. Peak decoding accuracy was obtained 150 ms after stimulus onset, compared to 110 and 80 ms after fixation onset (for F1 and F2 respectively; Figure 4b). Classifier weights during these peak-decoding latencies indicate the spatial distribution of category-differential information used for decoding (Figure 4c). The electrodes contributing to peak decoding locked to stimulus onset resemble the classic N170-effect topography (Figure 3b), while primarily occipital electrodes contribute to peak decoding of FRPs, resembling the topography of the FRP-lambda component (Figure 3b).

### Effects of baseline correction on decoding results

Finally, we turn to investigate the sustained category representation differences which yield prolonged above-chance decoding accuracy throughout the trial (Figure 3a and Figure 4a,c). Classifier weights in late time points within the trial show the qualitative contribution of occipital and temporal electrodes (Figure 4c). Traditionally, EEG data are baseline-corrected to remove slow drifts and assess activity differences evoked by the event of interest. However, if baseline periods contain systematic activity differences between conditions, baseline correction may induce these differences in later time points. This may cause misinterpretation of late effects within a trial, particularly noticeable in sustained decoding effects (van Driel et al., 2021). We re-analyzed category differences without baseline correction, extending the segment window size to -3500 to 3500 ms around stimulus onset, to include both the current and previous trials (We did not perform overlap correction over this entire window since we did not have enough data in this experiment for GLM convergence). Both univariate and multivariate analyses show an ongoing difference in category activity before and after stimulus onset (Figure 5a).

**Figure 5:**
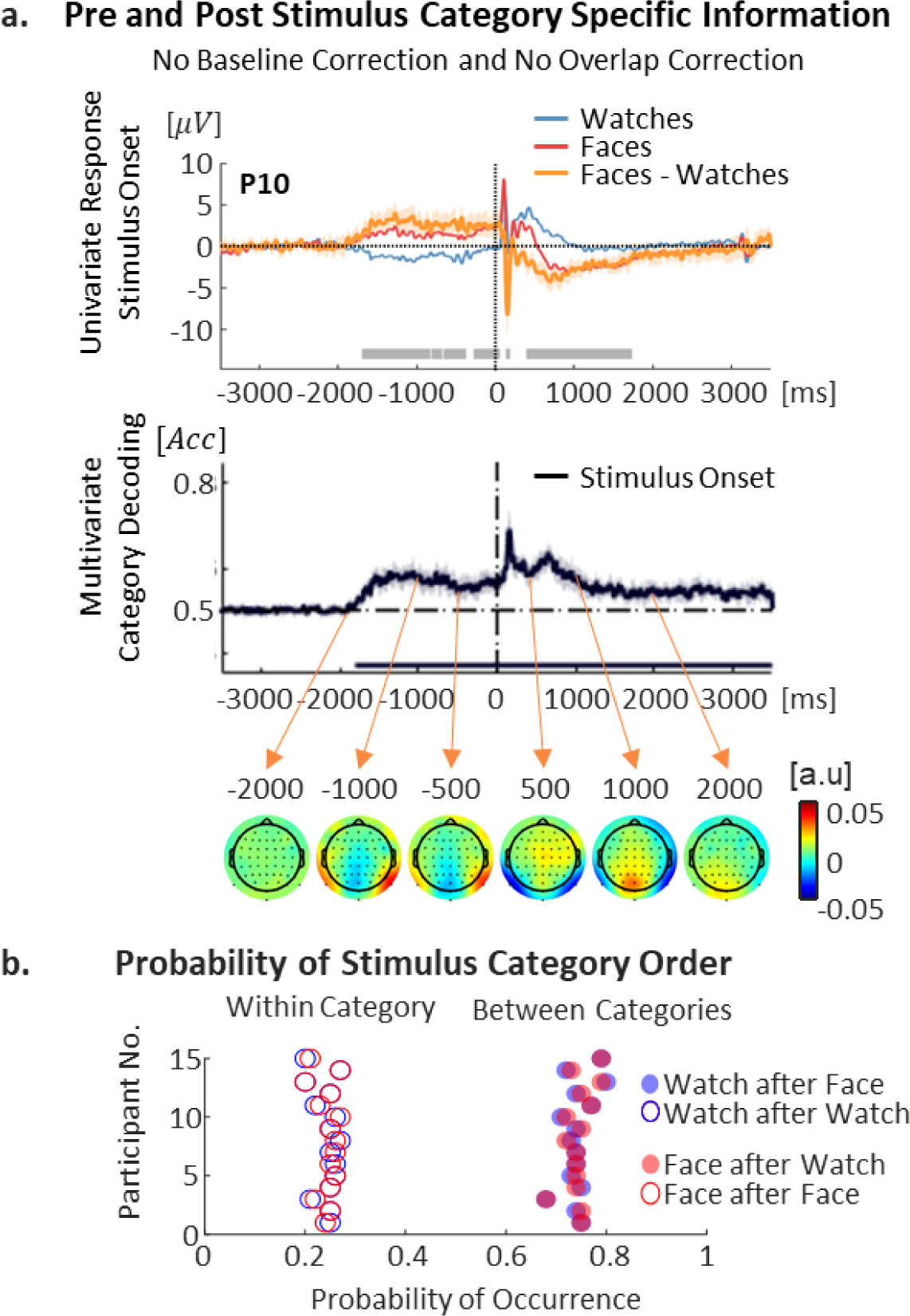
Pre-stimulus and post-stimulus category-specific information. **a.** Top: Grand average ERPs of raw data segments without baseline correction or overlap correction, time-locked to watch stimuli (blue), face stimuli (red), and the face-watch differential activity. Electrode P10 is presented during the window of -3500 to 3500 ms around stimulus onset, including the entire current and previous trials. The shaded intervals indicate 95% confidence intervals around the mean and the gray horizontal lines depict clusters in which the differential response is significantly different than zero using a two-sided cluster permutation test. Bottom: Time-resolved face vs. watch decoding accuracy from EEG segments time-locked to stimulus onset without baseline correction or overlap correction. The horizontal lines on the bottom depict time points during which the accuracy is significantly larger than chance using a permutation test. Topographies of the classifier weights are presented with arrows indicating the related time points within the segment. **b.** Probability of watch (blue) and face (red) stimuli to appear after a stimulus from the same category (empty circles) or a different category (full circles).

The topography of classifier weights indicates that the electrodes carrying category-specific information in the pre- and post-stimulus time points are reversed relative to the classifier separation plane. In ∼75% of trials, the stimulus category was different than the category of the stimulus preceding it (Figure 5b), since they were randomly chosen from a joint pool of faces and watches, with no effort in controlling the within-category and between-category stimuli pairs in the sequence. Faces were more likely to follow watches than other faces, and watches were more likely to follow faces than other watches (Figure 5b). It seems likely that this result reflects sustained category activation baseline differences between faces and watches in occipitotemporal electrodes, however, with this paradigm we cannot rule out pre-stimulus predictive activity.

## Discussion

In natural viewing conditions, observers typically make multiple fixations on objects of interest to gather detailed visual information. In this study we asked how object representations evolve across multiple fixations on the same object during free-viewing of prolonged stimuli. By combining fixation-related potentials (FRPs) and multivariate pattern analysis (MVPA), with methods to deconvolve temporally overlapping effects, we sought to determine if category-specific neural activity is re-evoked with each successive fixation and if sustained category representations can be tracked throughout the object’s presentation.

### Occipitotemporal Category Responses Are Evoked by Stimulus Onset, But Not Subsequent Fixations

Some studies suggest that eye movements counteract neural adaptation and perceptual fading in response to repeated or prolonged stimulation (Ditchburn and Ginsborg, 1952; Martinez-Conde et al., 2004). Shifting the gaze ensures a constant change of visual inputs, preserving response sensitivity to a stable environment, which could explain our sustained perception of objects throughout their presentation. However, this hypothesis would predict the reactivation of object representations with each fixation to support perceptual stability, which contradicts our findings. In our study, the N170 component, associated with category-specific structural encoding, was apparent when EEG activity was time-locked to stimulus onset, consistent with established literature (Bentin et al., 1996; Joyce and Rossion, 2005; Ganis et al., 2012; Rossion, 2014). However, this category-selective response was not re-evoked by subsequent fixations within the same object (Figure 3). Previous research has shown that fixation-related potentials, locked to the first fixation on an object during free viewing, show a face-selective N170 component captured by both univariate analyses and multivariate decoding (Auerbach-Asch et al., 2020, 2023; chapters 1 and 2 of this thesis). We thus conclude that the occipitotemporal category-selective response captured by the N170 is exclusive to the first acquisition of an object (whether passively or actively acquired) and does not recur with successive fixations. This result is in line with Spiering and Dimigen (2024) who concluded that face-emotion processing occurs upon stimulus onset but not following the firstmicrosaccades after ∼200 ms. However, their study was limited to very small saccades, during a fixation task, which creates minor changes in retinal input. The current study shows that even with larger saccades, of the magnitude elicited during natural viewing of faces from a conversational distance, category-selective activity is not re-ignited.

### Subsequent Fixations Evoke Early Occipital Responses

Using the GLM method to isolate the fixation-related response from the preceding stimulus-evoked response, we found that each subsequent fixation elicited an early (∼80-120 ms post-fixation) transient lambda component, thought to originate in the extrastriate cortex (Thickbroom et al., 1991; Shigeto et al., 1998; Kazai and Yagi, 2003; Rossion et al., 2003). Peak category-decoding accuracies based on occipital electrodes emerged within this same P1/lambda time window, likely reflecting differences in low-level visual processing attributes such as contrast, color, luminance, and spatial patterns (Kazai and Yagi, 1999; Picton et al., 2000; Tarkiainen et al., 2002; Nakashima et al., 2008). This aligns with MVPA findings showing exemplar-level decoding as early as 100 ms from stimulus onset, with higher-level category abstraction emerging later (Carlson et al., 2013; Cichy et al., 2014; Kaneshiro et al., 2015). Although we accounted for saccade amplitude and direction in the overlap deconvolution GLM, as well as the mean luminance and contrast, these parameters may not have been fully controlled for between different categories and may still contribute to the category decoding observed in the lambda component (Figure 2a).

### Multiple Fixations on an Object as a Form of Presentation Repetition

The absence of N170 reactivation across successive fixations may reflect a repetition suppression (RS) effect (Grill-Spector et al., 2006), as each fixation presents a slightly different view of the same object. RS is well-documented in response to repeated stimuli, where neural activity decreases due to mechanisms like local neural “fatigue”, altered response dynamics, or sparse neural coding (Kovács et al., 2006, 2013; Caharel et al., 2009; Eimer et al., 2010; Nemrodov and Itier, 2012; Feuerriegel et al., 2015). Adaptation effects tend to progressively increase along the visual hierarchy (Grill-Spector and Malach, 2001; Webster, 2015; Brands et al., 2024), potentially due to the increasing complexity of receptive fields and population response selectivity. Thus, we can consider sequential fixations within an object as a form of serial visual presentation (SVP), where each fixation shifts the object slightly on the retina, leading to adaptation in face-selective areas but less so in the early visual cortex, where receptive fields are smaller.

However, conceptualizing successive fixations as a form of SVP may overlook the active nature of information acquisition and integration across fixations. In natural vision, visual information is not only driven by bottom-up input at fixation onset but is also shaped by top-down signals, such as pre-saccadic parafoveal information and action-related sensory modulation, which provide prior context for the post-saccadic input (Huber-Huber et al., 2021). These signals enable the brain to form hypotheses about the underlying cause of the sensory activation (Friston et al., 2012; Parr and Friston, 2017). In SVP paradigms, prior information can typically be controlled, which is much harder to achieve in free-viewing contexts where participants actively acquire sensory input. From the perspective of predictive processing, brain activity encodes prediction errors between sensory input and internal models, with neural responses attenuated for expected inputs and heightened for unexpected ones (Summerfield and Egner, 2009; Friston, 2010). Although this framework applies to both SVP and free viewing, prior information tends to be richer in natural vision.

Real-world perception typically offers a pre-saccadic preview of post-saccadic information (Friston et al., 2012; Parr and Friston, 2017; Huber-Huber et al., 2021), and leads to face-selective N170 attenuation when a prediction of the saccade target object can be formed from peripheral cues (de Lissa et al., 2019; Huber-Huber et al., 2019; Buonocore et al., 2020). Using prolonged presentations of objects in our study, participants could obviously anticipate the basic category of the stimulus across eye movements (face or watch), and the transient occipital fixation-related response likely reflects low-level information processing that was not “explained away” by parafoveal predictions due to the coarse spatial resolution of peripheral vision. Our previous findings (Auerbach-Asch et al., 2020) indicate N170 attenuation during sequential fixations on separate exemplars of the same category, even though parafoveal information was limited and masked in that study, making pre-saccadic category expectations unlikely (see first chapter of this thesis). The ongoing debate about whether repetition suppression serves as a signature of prediction (Summerfield et al., 2008; Rostalski et al., 2020) or not (Tang et al., 2018; Feuerriegel, 2024), highlights the complexity of these processes, as both neural adaptation and predictive mechanisms may contribute to category-specific activity attenuation during free viewing observed in our study.

### Sustained object representation

After isolating the stimulus-evoked from fixation-evoked responses, we observed sustained category-specific information in occipitotemporal electrodes, time-locked to stimulus onset, that persisted beyond the transient onset response (>300 ms), as evidenced by prolonged above-chance decoding accuracies. Similar results were obtained using MEG in a study where participants maintained stable central fixation and avoided blinking during a 2-second object presentation (van de Nieuwenhuijzen et al., 2013). Supporting a sustained representation independent of refixations, intracranial ECoGcog studies investigating cortical responses to prolonged visual stimuli (Gerber et al., 2017; Vishne et al., 2023) have demonstrated sustained neural activation in the early visual cortex which track the duration of stimulus presentation even in the absence of saccadic eye movements. In contrast, more anterior and frontal regions respond primarily to stimulus onset, interpreted as reflecting onset-related processes such as categorization or object identification.

Category decoding accuracy peaked around 200 ms after stimulus onset, then decreased to a sustained plateau, accompanied by a reduction in the contribution of frontal electrodes to classification (Figure 3b and 4d). Although surface electrodes should not be confused with intracranial sources, we note that using intracranial recordings, Vishne et al. (2023) reported that occipitotemporal regions show sustained and stable category representations, while frontoparietal regions exhibit transient responses to stimulus onset. Further studies are needed to interpret the source of sustained category representations using EEG.

### Study limitations

The experimental paradigm in this study does not allow for a full dissociation of pre-stimulus expectations from post-stimulus sustained representations. In our design, category alterations (between faces and watches) occurred more frequently than category repetitions (Figure 5b), potentially allowing participants to predict the next stimulus. This predictability complicates the separation of sustained representations of the current stimulus from expectation effects related to the upcoming stimulus. Additionally, the interstimulus interval (ISI) of ∼400 ms was insufficient for EEG activity to fully return to baseline after stimulus offset. However, the above-mentioned MEG study (van de Nieuwenhuijzen et al., 2013) included a 2-second interstimulus interval (ISI) between stimuli, mitigating the risk of confusing prestimulus expectations with post-stimulus representations, which could be induced by baseline correction (see section *Effects of baseline correction on decoding results*). Thus, the results from both studies together suggest the presence of sustained category representations.

This study also emphasizes the methodological importance of signal overlap deconvolution of fixation-related potentials. Since our focus was on distinguishing evoked activity from successive fixations within a single object, it was crucial to separate each response from the preceding and subsequent events. Notably, decoding fixation-related activity without overlap correction resulted in above-chance decoding before fixation onset and sustained post-fixation decoding, indicating an overlap with the stimulus-evoked response (Figure 4a). However, the short ISIs between stimuli posed a challenge for deconvolving sustained effects, as slower overlapping responses require greater temporal jitter for successful deconvolution. For a detailed methodological discussion regarding signal overlap correction, readers are referred to (Dimigen et al., 2011; Nikolaev et al., 2016; Ehinger and Dimigen, 2019).

## Conclusions

This study examines how object representations unfold across successive fixations in natural viewing conditions. Category-specific information is extracted within 150 ms after an object appears, however subsequent fixations on the same object do not re-engage category-specific neural processes, as indicated by the absence of the N170 component. Instead, these fixations trigger early transient occipital responses, likely reflecting early visual cortex activation driven by low-level category differences. After deconvolving ensuing fixation-related potentials (FRPs), the stimulus-evoked response exhibits sustained category representations. Future research should aim to isolate pre-stimulus expectation effects from post-stimulus representations to clarify sustained neural dynamics in free-viewing conditions. Methodologically, this study highlights the importance of combining signal overlap deconvolution with multivariate analysis for accurate interpretation of neural activity during free-viewing. Taken together we hypothesize that exploratory fixations may serve as processing the “details” while simultaneously confirming the “gist” (Hochstein and Ahissar, 2002).

## Acknowledgments

This work was supported by grant 1902/14 from the Israel Science Foundation to L.Y.D. We thank all the research assistants for their help in data collection, and all the Deouell-lab members for their valuable comments and support.

## Notes

### Competing Interest Statement

The authors have declared no competing interest.

### Summary of Updates

I added a reference of a recent paper

